# Genetic Dissection of Mutual Interference between Two Consecutively Learned Tasks in *Drosophila*

**DOI:** 10.1101/2022.10.18.512721

**Authors:** Jianjian Zhao, Xuchen Zhang, Bohan Zhao, Liyuan Wang, Wantong Hu, Yi Zhong, Qian Li

## Abstract

Animals can continuously learn different tasks to adapt to changing environments and therefore have strategies to effectively cope with inter-task interference, including both proactive interference (Pro-I) and retroactive interference (Retro-I). Many biological mechanisms are known to contribute to learning, memory, and forgetting for a single task, however, mechanisms involved only when learning sequential different tasks are relatively poorly understood. Here, we dissect the respective molecular mechanisms of Pro-I and Retro-I between two consecutive associative learning tasks in *Drosophila*. Pro-I is more sensitive to inter-task interval (ITI) than Retro-I. They occur together at short ITI (<20 min), while only Retro-I remains significant at ITI beyond 20 min. Acutely overexpressing Corkscrew (CSW), an evolutionarily conserved protein tyrosine phosphatase SHP2, in mushroom body (MB) neurons reduces Pro-I, whereas acute knockdown of CSW exacerbates Pro-I. Such function of CSW is further found to rely on the γ subset of MB neurons and the downstream Raf/MAPK pathway. In contrast, manipulating CSW does not affect Retro-I as well as a single learning task. Interestingly, manipulation of Rac1, a molecule that regulates Retro-I, does not affect Pro-I. Thus, our findings suggest that learning different tasks consecutively triggers distinct molecular mechanisms to tune proactive and retroactive interference.

## Introduction

Continual learning is a natural ability of animals ranging from invertebrates to vertebrates but a great challenge for artificial intelligence (Fayek, Cavedon, & Wu, 2020; Kudithipudi et al., 2022; Parisi, Kemker, Part, Kanan, & Wermter, 2019; Wang et al., 2021). To achieve continual learning, the mutual interference between learning tasks, including proactive and retroactive interference, needs to be properly tuned. The interference from the previous task on the learning and memory of the current task is called proactive interference (Pro-I), while the interference from the current task on the memory of the following task is named retroactive interference (Retro-I) (Bouton, 1993; Wixted, 2004). Although we have much understanding of the molecular mechanisms of learning and memory for a single learning task (Johansen, Cain, Ostroff, & LeDoux, 2011; Kandel, Dudai, & Mayford, 2014), the molecular mechanisms that modulate the interferences between different tasks remain unclear.

*Drosophila* is a well-studied and highly tractable genetic model organism for understanding molecular mechanisms underlying a single learning task (Davis, 2005; Noyes, Phan, & Davis, 2021; Waddell & Quinn, 2001) as well as related human diseases (Mariano, Achsel, Bagni, & Kanellopoulos, 2020; Pandey & Nichols, 2011; van Alphen & van Swinderen, 2013). In recent years, *Drosophila* has also emerged as an excellent model for studying interference mechanisms between two associative learning tasks. Several molecules, including Rac1, Foraging, Scribble, SLC22A, Fmr1, SCAR and Dia, have been reported to regulate Retro-I (Cervantes-Sandoval, Chakraborty, MacMullen, & Davis, 2016; Dong et al., 2016; Gai, Liu, Cervantes-Sandoval, & Davis, 2016; Gao et al., 2019; Reaume, Sokolowski, & Mery, 2011; Shuai et al., 2010). Of these molecules, Scribble and SLC22A, have also been reported to regulate Pro-I (Cervantes-Sandoval et al., 2016; Gai et al., 2016). It indicates that Retro-I and Pro-I may have shared regulatory molecules. Previous studies reported that resistance to Pro-I and Retro-I is very different in patients with autism (Mottron, Belleville, Stip, & Morasse, 1998), schizophrenia (Torres, Flashman, O’Leary, & Andreasen, 2001), and ADHD (Orban, Festini, Yuen, & Friedman, 2022). It raises a possibility that Pro-I and Retro-I may have their distinct molecular mechanisms. Of note, all reported molecules in regulating Retro-I or Pro-I also play important roles in the learning or memory of a single learning task (Cervantes-Sandoval et al., 2016; Dong et al., 2016; Gai et al., 2016; Gao et al., 2019; Reaume et al., 2011; Shuai et al., 2010). Whether there are molecules specifically responsible for modulating memory interference without affecting a single learning task remains to be determined.

## Results

### Pro-I and Retro-I differ in time interval sensitivity and regulatory molecules

Consistent with previous studies (Cervantes-Sandoval et al., 2016; Gai et al., 2016), associative learning as a proactive experience caused Pro-I to the target task in wild-type flies (Canton-S) (Figure 1-figure supplement 1). In studies of Retro-I, memory for the target task is significantly impaired regardless of whether the retroactive experience is associative learning (Cervantes-Sandoval et al., 2016; Dong et al., 2016; Gai et al., 2016; Gao et al., 2019; Reaume et al., 2011; Shuai et al., 2010) or non-associative stimuli (Mo et al., 2022; Zhang, Li, Wang, Liu, & Zhong, 2018). Unlike Retro-I, non-associative stimuli (Odor Alone, Shock Alone, and Backward group) did not affect the learning of the target task when used as a proactive experience compared to the Naïve group (Figure 1-figure supplement 1).

Retro-I can be observed regardless of whether the time interval between two tasks is short or long (Cervantes-Sandoval et al., 2016; Dong et al., 2016; Gai et al., 2016; Gao et al., 2019; Reaume et al., 2011; Shuai et al., 2010). We tested the relationship between the Pro-I and inter-task interval (ITI) (Figures 1A-1C). Flies showed significant Pro-I when the ITI was 15 min or less (0, 5, 10, and 15 min). If the ITI was 20 min or more (20, 30, and 60 min), no significant Pro-I was observed. In contrast, when the ITI was gradually increased from 0 to 60 min (0, 20, 30, and 60 min), the flies consistently exhibited significant Retro-I (Figures 1D and 1E).

**Figure 1.**
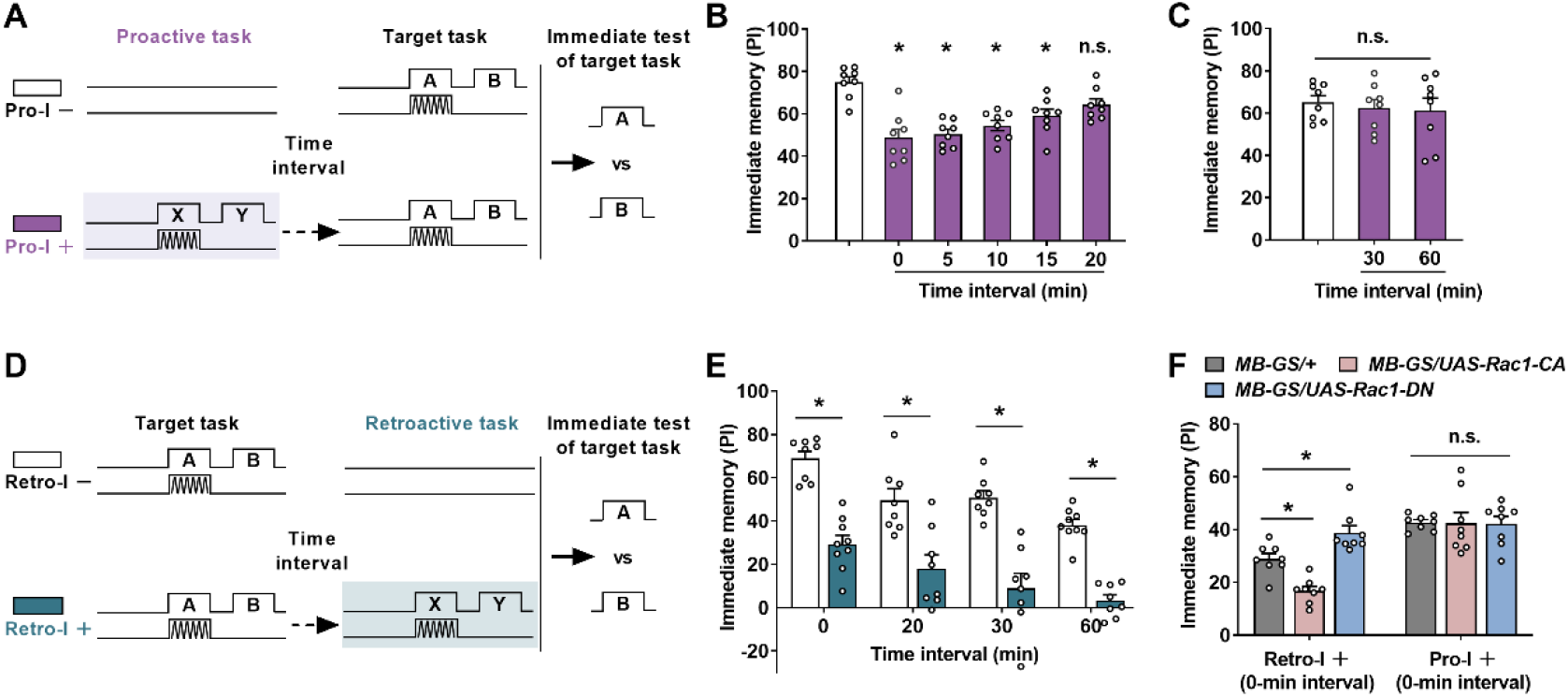
Differences between Pro-I and Retro-I. (A-C) The paradigm (A) and the behavioral results (B and C) of the proactive interference (Pro-I) experiment. The time interval between the proactive task and the target task was changed from 0 min to 60 min. Proactive interference was significant when the time interval was less than 20 min (0 min, 5 min, 10 min, or 15 min) in wild-type flies (middle and right). n = 8 (D and E) The paradigm (D) and the behavioral results (E) of the retroactive interference (Retro-I) experiment. Retroactive interference was significant when the time interval between the target and the retroactive task was 0 min, 20 min, 30 min, and 60 min. n = 8-9. (F) The behavioral performance of transgenic flies with retroactive or proactive interference. Compared to the control group (*MB-GS/*+, RU486+), Rac1-CA-expressing flies (*MB-GS/UAS-Racl-CA*, RU486+) showed a significantly lower performance index, while Rac1-DN-expressing flies (*MB-GS/UAS-Racl-DN*, RU486+) exhibited a higher memory index in retroactive interference (Retro-I). No significant difference was observed in all groups with proactive interference (Pro-I). n = 8. Statistics: ordinary one-way ANOVA with Dunnett’s multiple comparisons test (B and C); two-way ANOVA with Bonferroni’s multiple comparisons test (E and F). Results with error bars are means ± SEM. *p < 0.05. n.s., non-significant.

Given that Scribble and SLC22A regulate both Pro-I and Retro-I (Cervantes-Sandoval et al., 2016; Gai et al., 2016), other regulators of Retro-I may also affect Pro-I. Rac1 is required to mediate Retro-I when the time interval between two tasks is 1.5 h (Shuai et al., 2010). We next tested whether Rac1 also affects Retro-I with 0-min ITI and explored whether Rac1 plays a role in Pro-I (Figure 1F). The transgene dominant-negative Rac1 (Rac1-DN) or constitutively active Rac1 (Rac1-CA) was used to inhibit or increase Rac1 activity (Luo, Liao, Jan, & Jan, 1994). Rac1-DN or Rac1-CA was expressed using MB-GS, which is an inducible Gene-Switch (GS) driver for mushroom body (MB) neurons used to allow transgene expression only on administration of the drug RU486 (Mao, Roman, Zong, & Davis, 2004). Suppressing (*MB-GS/UAS-Rac1-DN*, RU486+) or increasing (*MB-GS/UAS-Rac1-CA*, RU486+) Rac1 activity in MB neurons significantly mitigated or aggravated the Retro-I. However, the same manipulations did not affect the Pro-I, indicating that there is a different molecular mechanism underlying the Pro-I.

### Pro-I, but not Retro-I, is bidirectionally regulated by CSW

Since the difference between Pro-I and Retro-I lies in the sensitivity to inter-task time intervals, we speculate that molecules that modulate “time interval” effects may specifically regulate Pro-I. Given that CSW, an evolutionarily conserved protein tyrosine phosphatase SHP2, has been reported to regulate the “time interval” effect in long-term memory (LTM) formation in *Drosophila* (Pagani, Oishi, Gelb, & Zhong, 2009), we tested the role of CSW in Pro-I. Compared to the parental control group (*MB-GS/+*, RU486+), acutely knocking down CSW in MB neurons using two independent RNAi lines (*MB-GS/UAS-csw-RNAi-1* and *MB-GS/UAS-csw-RNAi-2;* RU486+) showed more severe Pro-I (Figure 2A), which can be better reflected by the Pro-I+/Pro-I– ratio (Figure 2B). To further determine this result, we added a comparison with uninduced control groups (RU486–) and obtained consistent results (Figure 2C). When the interval between the two tasks was 20 min, Pro-I was no longer observed in control flies, but was still present in flies with acute knockdown of CSW (Figure 2D). Conversely, acute overexpression of CSW in MB neurons (*MB-GS/UAS-csw*, RU486+) reduced Pro-I by using a newly constructed transgenic strain (Figure 2E and 2F) and a previously reported strain (Botham, Wandler, & Guillemin, 2008) (Figure 2-figure supplement 1). Of note, bidirectional manipulation of CSW expression in MB neurons did not affect Retro-I (Figure 3G).

**Figure 2.**
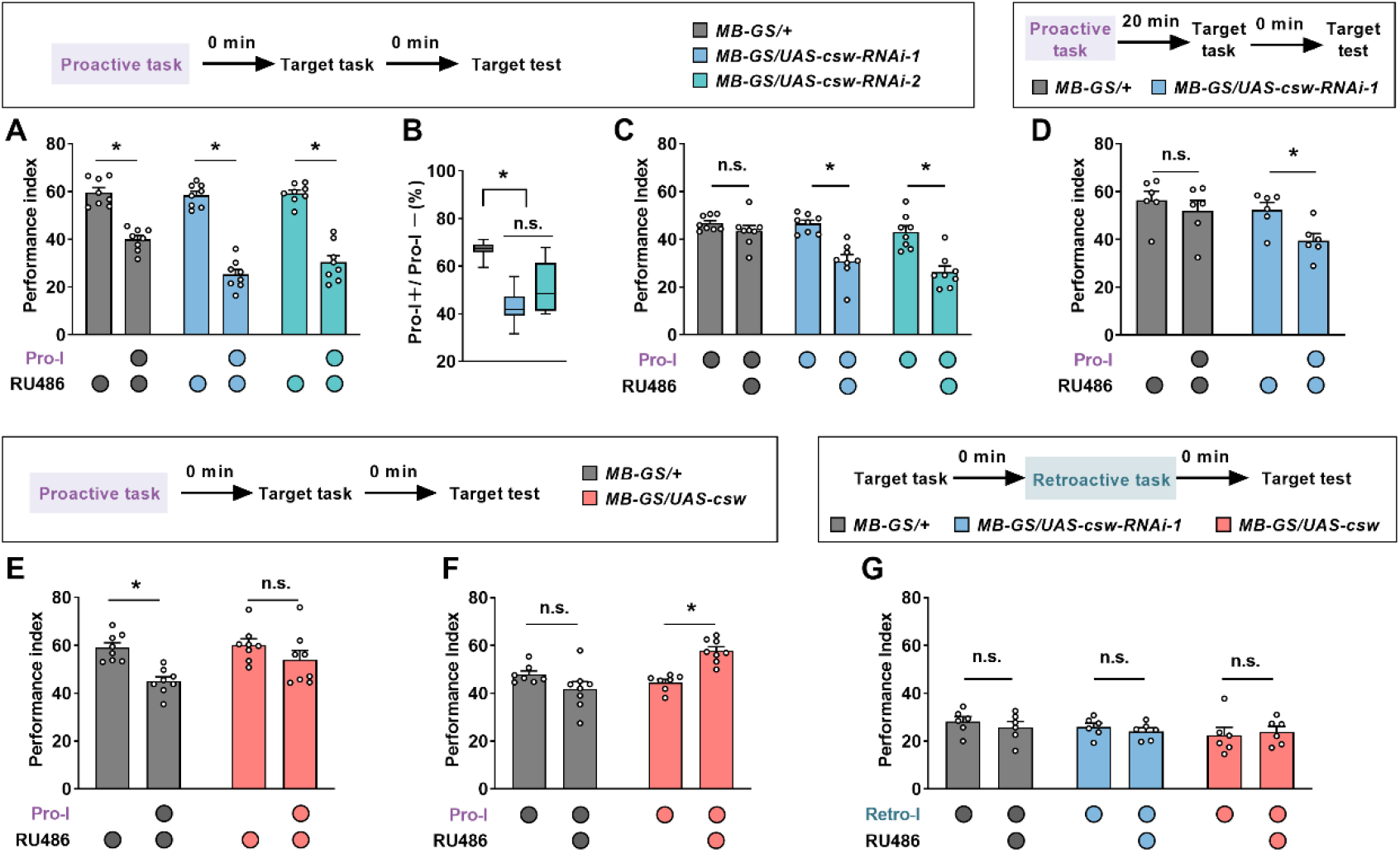
CSW Bidirectionally Regulates Proactive but not Retroactive Interference. (A-C) The behavioral performance of transgenic flies with proactive interference (0-min interval). Acute knockdown of CSW in mushroom body neurons (*MB-GS/UAS-csw-RNAi-1* or *MB-GS/UAS-csw-RNAi-2*; RU486+) led to more severe proactive interference relative to the genetic control group (*MB-GS/+*, RU486+) (A and B) and uninduced controls (RU486-) (C). n = 8. (D) The behavioral performance of transgenic flies with proactive interference (20-min interval). csw-RNAi-expressing flies (*MB-GS/UAS-csw-RNAi-1*, RU486+) but not control flies (*MB-GS/+*, RU486+) showed significant proactive interference. n = 6. (E and F) The behavioral performance of transgenic flies with proactive interference (0-min interval). Acute overexpression of CSW in mushroom body neurons (*MB-GS/UAS-csw*; RU486+) prevented proactive interference compared with the genetic control group (*MB-GS/+*, RU486+) (E) and uninduced controls (RU486-) (F). n = 8. (G) The behavioral performance of transgenic flies with retroactive interference (0-min interval). Acute knockdown (*MB-GS/UAS-csw-RNAi-1*, RU486+) or overexpression (*MB-GS/UAS-csw*, RU486+) of CSW in mushroom body neurons did not affect retroactive interference compared with genetic and uninduced controls. n = 6. Statistics: ordinary one-way ANOVA with Dunnett’s multiple comparisons test (B); two-way ANOVA with Bonferroni’s multiple comparisons test (A, and C-G). Results with error bars are means ± SEM. *p < 0.05. n.s., non-significant.

**Figure 3.**
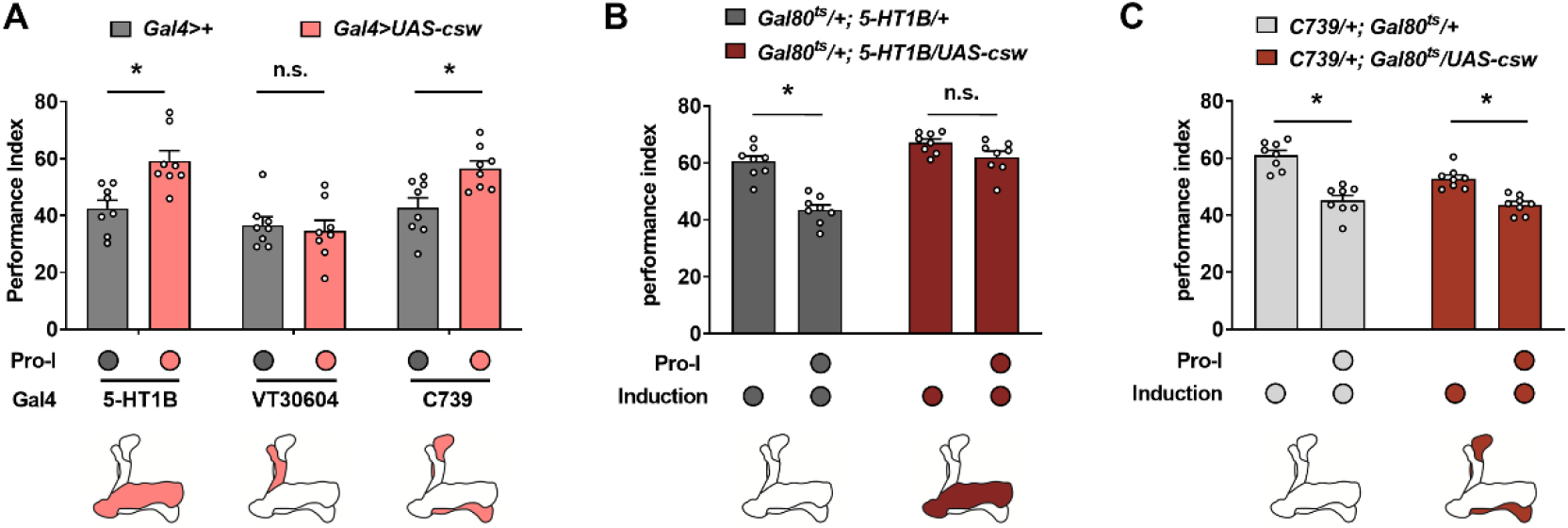
Acutely Overexpressing CSW in MB γ Neurons Prevents Proactive Interference. The immediate memory performance with or without proactive interference (Pro-I, 0-min interval) was tested in different transgenic flies (A-C). (A) Flies with overexpression of CSW in MB γ neurons (*5-HT1B*>*UAS-csw*) and α/β neurons (*C739>UAS-csw*) showed higher memory performance than their genetic control group. n = 8. (B) Acutely overexpressing CSW in MB γ neurons (*Gal80^ts^/+; 5-HT1B/UAS-csw*) prevented proactive interference relative to the genetic control group (*Gal80^ts^/+; 5-HT1B/+*). n = 8. (C)Acutely overexpressing CSW in MB α/β neurons (*C739/+; Gal80^ts^/UAS-csw*) showed similar proactive interference with the genetic control group (*C739/+; Gal80^ts^/+*). n = 8. Statistics: two-way ANOVA with Bonferroni’s multiple comparisons test. Results with error bars are means ± SEM. *p < 0.05. n.s., non-significant.

### Pro-I is reduced by overexpressing CSW in MB γ Neurons

The MB contains ~2,000 intrinsic neurons called Kenyon cells (KCs), which are further divided into three major types: γ neurons (~675), α/β neurons (~990), and α’/β’neurons (~350) (Aso et al., 2014). We next sought to determine whether the modulatory effects of CSW on Pro-I could be narrowed to a certain type of MB neurons. 5-HT1B-Gal4 (Gao et al., 2019; Shyu et al., 2017; Yuan, Lin, Zheng, & Sehgal, 2005), VT30604-Gal4 (Wu, Shih, Lee, & Chiang, 2013), and C739-Gal4 (O’Dell, Armstrong, Yang, & Kaiser, 1995) were used to drive transgene expression in γ neurons, α’/β’ neurons, and α/β neurons. Overexpressing CSW in γ neurons and α/β neurons, but not in α’/β’ neurons showed higher performance with Pro-I when compared with their respective genetic controls (Figure 3A). To rule out possible developmental effects, we employed another inducible expression system called TARGET, which relies on a temperature shift to induce transgene expression (McGuire, Le, Osborn, Matsumoto, & Davis, 2003). Acute overexpression of CSW in γ neurons (*Gal80^ts^/+; 5-HT1B/UAS-csw*, induced), but not in the α/β neurons (*C739/+; Gal80^ts^/UAS-csw*, induced), significantly reduced the Pro-I compared with their respective genetic controls (Figure 4B and 4C). These data suggest that CSW regulates Pro-I in MB γ neurons.

**Figure 4.**
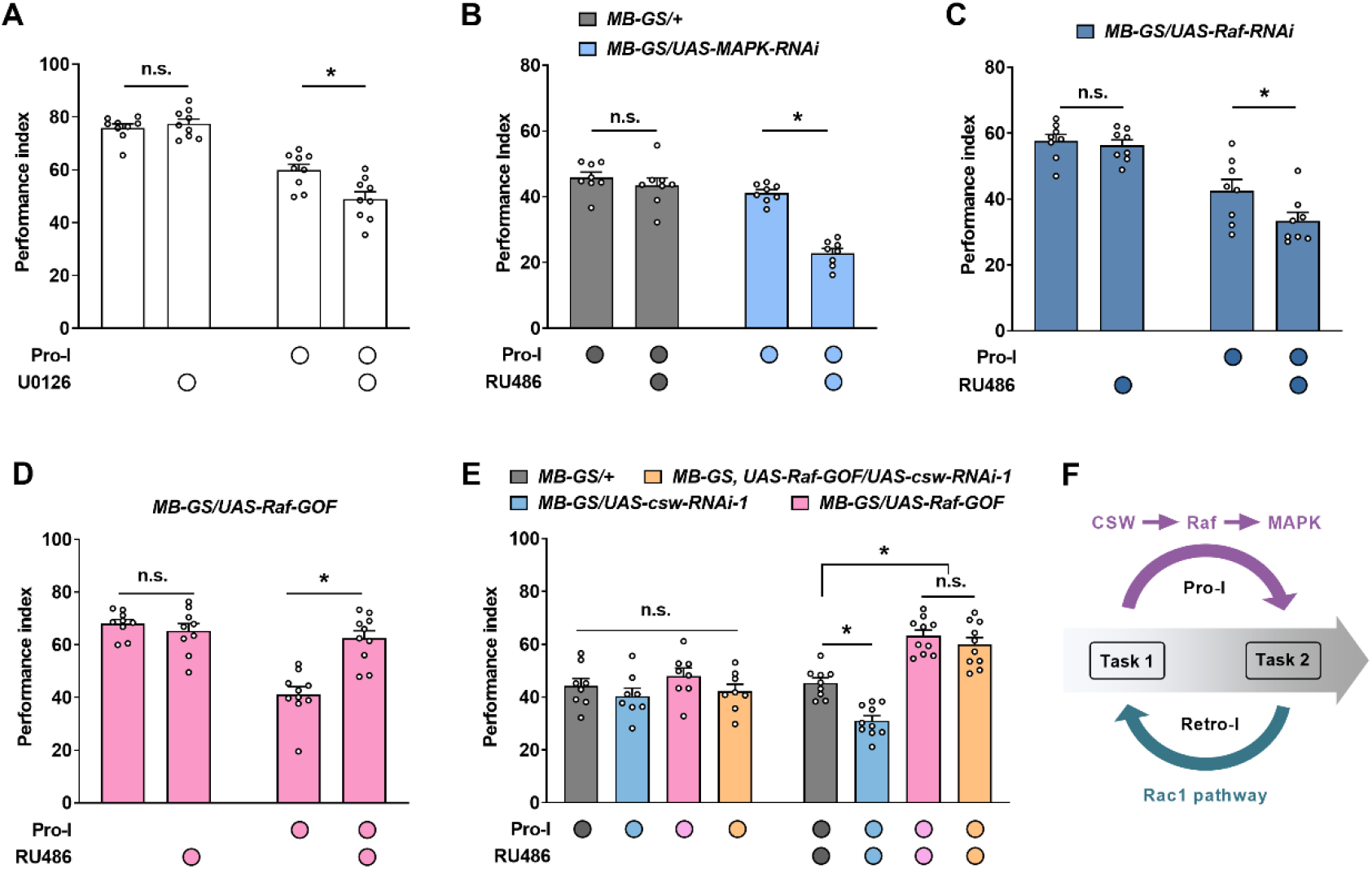
CSW Regulates Proactive Interference through Raf/MAPK pathway. The immediate memory performance with or without proactive interference (Pro-I, 0-min interval) was tested in different transgenic flies (A-E). (A) Pharmacological inhibition of MAPK by feeding U0126 inhibitor aggravated the proactive interference in wild-type flies. n = 8. (B) Flies with acute knockdown of MAPK in MB neurons (MB-GS/UAS-MAPK-RNAi, RU486+) exhibited more severe proactive interference compared with genetic and uninduced controls. n = 8. (C) Acute knockdown of Raf in MB neurons (*MB-GS/UAS-Raf-RNAi*, RU486+) aggravated the proactive interference relative to the uninduced control. n = 8. (D) Acutely overexpressing Raf-GOF in MB neurons (*MB-GS/UAS-Raf-GOF*, RU486+) reduced the proactive interference compared with its uninduced control group. n = 8. (E) Acute overexpression of Raf-GOF (*MB-GS/UAS-Raf-GOF*, RU486+) dominated the effect of CSW knockdown (*MB-GS/UAS-csw-RNAi*, RU486+) on proactive interference. No significant difference was found between uninduced groups without RU486 feeding. n = 10. (F) Model of molecular mechanisms underlying proactive and retroactive interference. Statistics: two-way ANOVA with Bonferroni’s multiple comparisons test. Results with error bars are means ± SEM. *p < 0.05. n.s., non-significant.

### The Raf/MAPK pathway acts downstream of the CSW to regulate Pro-I

CSW is generally considered as a positive regulator of Ras/MAPK signaling (Perkins, Johnson, Melnick, & Perrimon, 1996). And the regulation of CSW on the “time interval” effect in LTM formation is also thought to be via the MAPK pathway (Pagani et al., 2009). So we further explored whether CSW affects Pro-I through the MAPK pathway. Feeding U0126, a widely used inhibitor of the MAPK pathway (Thomas & Huganir, 2004), did not affect single-task learning, but significantly exacerbated Pro-I when learning two consecutive tasks (Figure 4A). Flies with acute genetic knockdown of MAPK in MB neurons (*MB-GS/UAS-MAPK-RNAi*, RU486+) also exhibited more severe Pro-I than uninduced and parental control flies (Figure 4B).

Consistent with these results, Raf kinase, a classical upstream regulator of MAPK (Thomas & Huganir, 2004), was found to bidirectionally regulate the Pro-I in MB neurons like CSW (Figures 4C and 4D). Acutely knocking down Raf significantly exacerbated the Pro-I, while acute overexpression of Raf-GOF, which encodes a constitutively active Raf kinase (Brand & Perrimon, 1994), significantly reduced Pro-I. When Raf-GOF and CSW-RNAi were co-expressed, Raf-GOF expression dominated the effect on Pro-I, indicating that Raf acts downstream of CSW (Figure 5E). Thus, our data support that CSW regulates Pro-I through Raf/MAPK pathway.

Our previous study found that the Raf/MAPK pathway is activated by learning to protect memory retention via non-muscle myosin II Sqh after single-task learning (Zhang et al., 2018). Although the regulatory effect of CSW on Pro-I in two consecutive task learning was also through the Raf/MAPK pathway, manipulating CSW did not affect the learning and memory of a single task (Figure 4-figure supplement 1A and 1B), unlike Raf (Figure 4-figure supplement 1C) (Zhang et al., 2018). Interestingly, as a downstream molecule of the Raf/MAPK pathway in regulating single-task memory (Zhang et al., 2018), Sqh did not participate in the Pro-I between the two tasks (Figure 4-figure supplement 1D). These results suggest that learning two different tasks consecutively may specifically recruit CSW to activate the Raf/MAPK pathway to reduce Pro-I through downstream molecules different from Sqh, a mechanism that differs from the memory protection mechanism triggered by a single learning task.

## Discussion

Our findings support a model for understanding molecular mechanisms of mutual interference between two successive learning tasks in *Drosophila* (Figure 4F). First, in learning two different associative tasks in succession, Pro-I and Retro-I occur simultaneously when the time distance between tasks is close (< 20 min), while only Retro-I could be observed when the time distance was far (> 20 min). Second, we identify a molecular pathway that specifically regulates Pro-I: the CSW/Raf/MAPK pathway. The upstream signaling molecule CSW of this pathway is not involved in single-task learning and memory. In learning two different tasks close in time, although Pro-I and Retro-I occurred simultaneously, CSW was only involved in regulating Pro-I. Third, Rac1 specifically regulates Retro-I without affecting Pro-I when two different tasks are temporally close. Consistently, the Rac1/SCAR/Dia pathway has been reported to regulate Retro-I when the two tasks are temporally distant (1.5-h interval) (Gao et al., 2019; Shuai et al., 2010).

In the mammalian brain, sensory information is reported to converge to associative brain regions, such as the prefrontal cortex and medial temporal lobe (Markov et al., 2013). Such convergence is thought to be important for flexibility in working memory, but can also cause capacity limitations due to interference between items in memory, suggesting that there is a trade-off between flexibility and interference resistance (Buschman, 2021). In *Drosophila*, sensory information from different modalities can be transmitted to the MB, the key site of associative learning, and converges to a small set of output neurons (MBONs) to guide different behaviors (Li et al., 2020; Modi, Shuai, & Turner, 2020). This convergence may also give rise to a trade-off between flexibility and interference resistance of associative memory. Consistent with this idea, acute inhibition of Rac1 or Fmr1 mutants are resistant to Retro-I but exhibit defects in behavioral flexibility reflected by reversal learning (Dong et al., 2016; Shuai et al., 2010). Knocking down Scribble was able to decrease both Retro-I and Pro-I but also displayed a defect in reversal learning (Cervantes-Sandoval et al., 2016). We show that Pro-I regulated by the CSW/Raf/MAPK pathway differs from Rac1-mediated Retro-I. It is interesting to test whether CSW is also involved in regulating behavioral flexibility. Although Pro-I in the current work only occurs when the interval between two tasks is less than 20 min, the phenomenon of Pro-I varies across biological learning systems and tasks (Crossley et al., 2019; Epp, Mera, Kohler, Josselyn, & Frankland, 2016; Jonides & Nee, 2006). Future works are required to determine whether orthologues of CSW also participate in other forms of Pro-I. Interestingly, the molecules regulating Pro-I and Retro-I in *Drosophila* are involved in different diseases with varying levels of intellectual disability. Rac1 and Fmr1, which regulate Retro-I in *Drosophila*, are high-risk genes for autism (Lord et al., 2020). Mutations in *PTPN11*, a human orthologue of *csw*, account for more than half of Noonan syndrome (Roberts, Allanson, Tartaglia, & Gelb, 2013). Therefore, further studies on the molecular mechanisms underlying Pro-I and Retro-I may also contribute to the understanding of the pathogenesis of autism and Noonan syndrome.

The spacing effect of memory was discovered more than a century ago: repeated learning at long time intervals produces significantly better memory performance than repeated learning at short or no time intervals (Ebbinghaus, 1885/1913; Smolen, Zhang, & Byrne, 2016). The optimal time interval for repeated learning in this spacing effect has been found to correlate with the peak time of MAPK activation triggered by each learning (Pagani et al., 2009; Philips, Tzvetkova, & Carew, 2007; Philips, Ye, Kopec, & Carew, 2013). Overexpression of CSW has been reported to shorten the peak time of learning-activated MAPK, thus allowing repeated learning at short intervals (45 s) to produce superior memory performance that is dependent on long intervals (15 min) in *Drosophila* (Pagani et al., 2009). In the current study, we show that the time interval between two successive different tasks determines the presence of Pro-I. Overexpression of CSW reduces Pro-I between two consecutive different tasks (0-min interval), achieving the same effect that is only available when the inter-task interval is longer than 20 min. Together with previous studies (Pagani et al., 2009), our data suggest that the modulation of spacing effect and Pro-I by CSW occurs in different groups of neurons. Overexpression of CSW in MB α/β neurons is sufficient to regulate the spacing effect in repeated learning of the same task (Pagani et al., 2009), while overexpression of CSW in MB γ neurons affects Pro-I between two different tasks. A common feature of these two distinct functions of CSW is the recruitment of the MAPK pathway for the regulation of the time interval between learning events. Although a learning event can physically occur quickly, its biological representation may take more time to complete, depending on a molecular context established by MAPK (Philips et al., 2013). With this idea, a possible unifying explanation is that CSW can accelerate the construction of MAPK contexts and thus obtain a more complete biological representation of a learning event in a shorter time. For the spacing effect, longer time intervals allow the same learning task to be repeated more completely and therefore memory is better formed. For Pro-I, if the proactive task does not have enough time to be fully represented in the MAPK context, it may take up resources for learning a subsequent target task and thus cause interference.

## Materials and Methods

### Fly Stocks

Flies (*Drosophila melanogaster*) were cultured at 23 °C and 60% relative humidity with standard medium under a 12 h light-dark cycle. Flies using the TARGET system were raised at 18 °C. *MB-GS* was a gift from Dr. Ronald L. Davis (Mao et al., 2004). *VT30604-Gal4* was a gift from Dr. Ann-Shyn Chiang (Wu et al., 2013). *Canton-S* (#64349), *UAS-Rac1-CA* (#6291), *UAS-Rac1-DN* (#6292), *UAS-csw* (#23878), *UAS-csw-RNAi-1* (#31760), *UAS-csw-RNAi-2* (#33619), *UAS-Raf-GOF* (#2033), *UAS-MAPK-RNAi* (#31524), *5-HT1B-Gal4* (#27637), and *Gal80^ts^* (#7019 and #7017) were obtained from Bloomington Stock Center. The *UAS-Raf-RNAi* (#5796) and *UAS-sqh-RNAi* (#1223) were acquired from Tsinghua Fly Center. The *C739-Gal4* (O’Dell et al., 1995) was the extant stock in our lab.

### Generation of transgenic flies

Since the insertion site of *UAS-csw* in the transgenic line (#23878) acquired from Bloomington Storck Center is the X chromosome, a distinction between male and female flies needs to be made when counting behavioral results. For experimental convenience, we constructed a new transgenic strain (*UAS-csw*, insertion on chromosome III) and used it mainly in this study. Construction of this strain was performed at Fungene Biotech (http://www.fungene.tech). NotI/XbaI PCR fragment of coding sequences of *csw-RA* was inserted into the NotI/XbaI sites of pJFRC28-10XUAS-IVS-GFP-p10 vector (Addgene Plasmid #36431), and then the construct was inserted into *attp2* site.

### Behavioral Assays

Flies from 2 to 5 days old were reared for behavioral experiments under the Pavlovian olfactory conditioning procedure. (Tully, Preat, Boynton, & Delvecchio, 1994; Tully & Quinn, 1985). The flies were first transferred to a behavior room at 23°C and 60% relative humidity for 30 min to adapt. The training time for each associative task was 5 min. About 100 flies were subjected to 90 s of air, 60 s of odor exposure accompanied by 12 pulses of electric shock at 60V (conditioned stimulus +, CS+), 45 s of air, 60 s of another different odor (conditioned stimulus –, CS–) and 45 s of air. To test memory, trained flies were placed in a T-maze to choose between two odors, CS+ and CS–. After 1 min of the choice, the performance index (PI) can be calculated based on the distribution of the flies between the two odors. A PI of 100 indicates that all flies escape CS+ odor, while a PI of 0 means that flies have no preference for CS+ and CS–. OCT (3-octanol, 1.5×10^-3^ in dilution, Aldrich) and MCH (4-methylcyclohexanol, 1.0×10^-3^ in dilution, Fluka) were used for learning and testing of the target task in Pro-I and Retro-I paradigm. To balance naive odor bias, OCT and MCH were used as CS+ and CS– reciprocally, and the complete performance index was defined as the average. EA (ethyl acetate, 2×10^-3^ in dilution, Alfa Aesar) and IA (isoamyl acetate, 2×10^-3^ in dilution, Alfa Aesar) were used as CS+ and CS– for learning of the non-target task (proactive or retroactive).

### Drug Feeding Treatment

Flies were fed with drugs as previously described (Zhang et al., 2018). Control flies (RU486– or U0126–) were fed a control solution containing 5% glucose and 3% ethanol. Flies were fed with 500 μM RU486 (Mifepristone, J&K) dissolved in a control solution in RU486+ groups. Flies were fed with 20 μM U0126 (Cell Signaling Technology) dissolved in a control solution for 16 h before training.

### Transgene Induction

In this study, two inducible systems, GeneSwitch (Mao et al., 2004) and TARGET (McGuire et al., 2003), were used for transgene expression. In the GeneSwitch system, the transgene expression was induced by RU486 feeding for two days. In the TARGET system, flies that were raised at 18°C were transferred to a 31°C incubator for 3 days to induce the transgene expression.

### Statistics

Statistical analysis was performed using Prism (Graphpad). Four normality tests were used, including Anderson-Darling, D’Agostino-Pearson omnibus, Shapiro-Wilk, and Kolmogorov-Smirnov. All experimental data passed at least two normality tests. Comparisons of multiple groups were performed using ordinary one-way ANOVA with Dunnett’s multiple comparisons test or two-way ANOVA with Bonferroni’s multiple comparisons test. P values < 0.05 were considered statistically significant and marked with *, and n.s. means non-significant differences (p > 0.05).

## Acknowledgments

We thank Dr. Ronald L. Davis, Dr. Ann-Shyn Chiang, and the Bloomington Stock Center for fly stocks. This work was supported by grants from the National Natural Science Foundation of China (31970955, to Q.L.; 32021002, to Yi Zhong) and the Tsinghua-Peking Center for Life Sciences.

## Author contributions

Q. L., X. Z., and B. Z. contributed to the initial idea of this project. Q.L. and Y. Z. supervised this project. Q. L. and Z. J. designed all experiments. J. Z. performed experiments and analyzed the results. J. Z., W. H., L. W., and Q. L. contributed to the writing of the manuscript.

---

**Figure 1-figure supplement 1.**
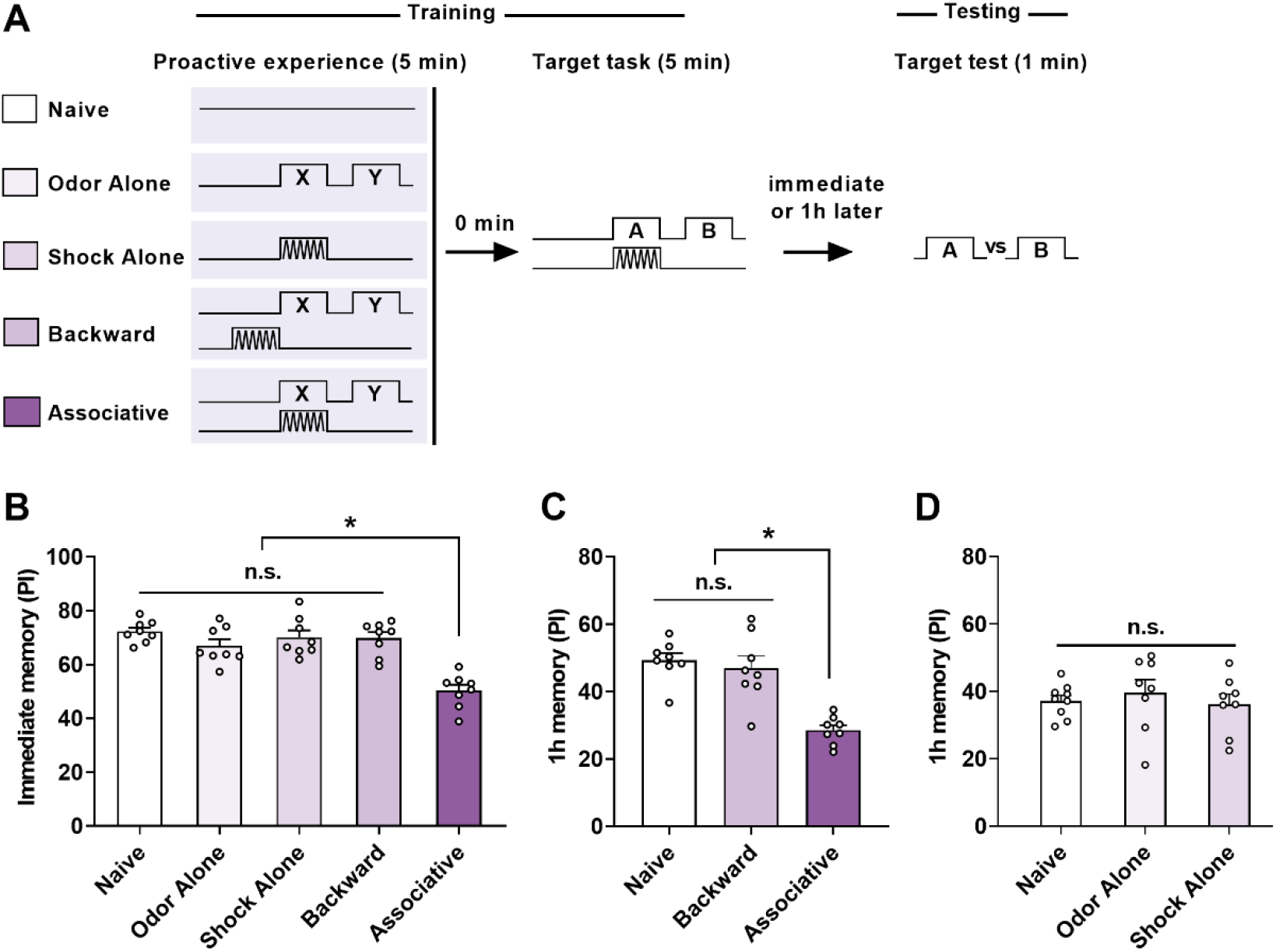
Effect of Associative and Non-associative Factors on Proactive Interference. (A) Behavioral schedules. A naïve group, three non-associative control groups (Odor Alone, Shock Alone, and Backward), and an associative group (Associative) were used as proactive experiences. (B) Comparison of immediate memory performance of the target task. Compared with the Naïve group, significant proactive interference was observed in the Associative group, but not in non-associative control groups (n = 8). (C) Comparison of 1-h memory performance of the target task. Compared with the Naïve group, significant proactive interference was observed in the Associative group, but not in the Backward control group. n = 8. (D) Comparison of 1-h memory performance of the target task. No significant difference was found between Naïve, Odor Alone, and Shock Alone groups. n = 8-9. Statistics: ordinary one-way ANOVA with Dunnett’s multiple comparisons test. Results with error bars are means ± SEM. *p < 0.05. n.s., non-significant.

**Figure 2-figure supplement 1.**
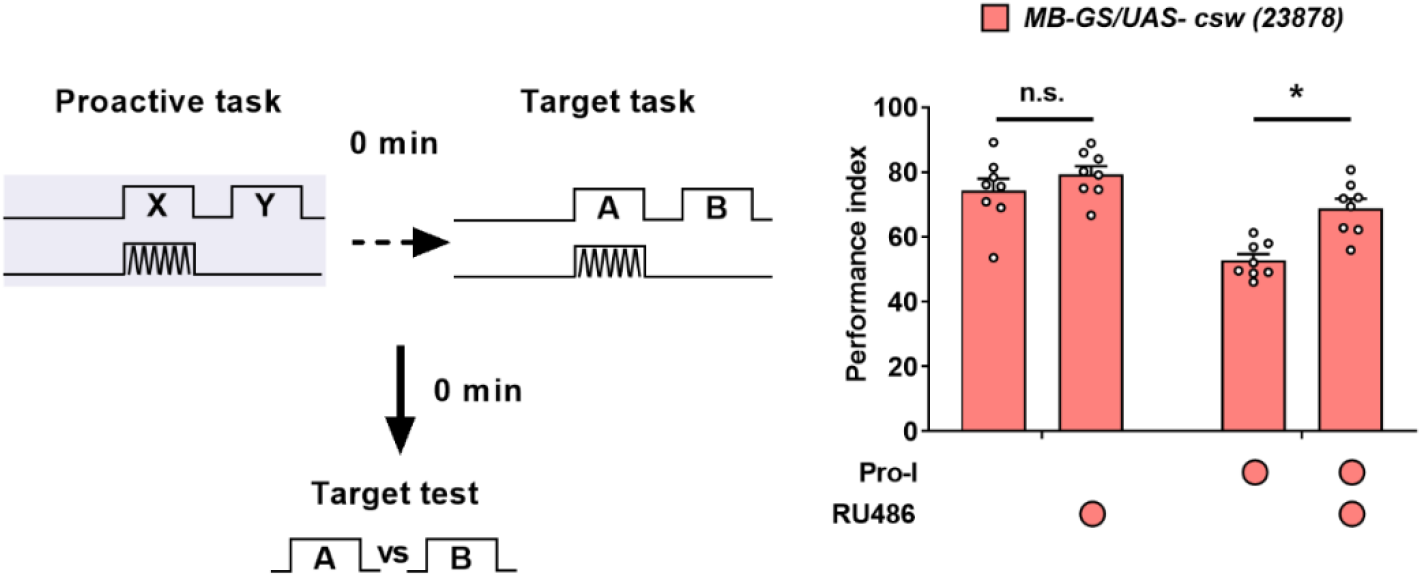
Overexpressing CSW in MB neurons reduces proactive interference. The paradigm (left panel) and the behavioral results (right panel) of the proactive interference (Pro-I) experiment. Acutely overexpressing CSW in MB neurons (*MB-GS/UAS-csw (23878)*, RU486+) reduced the proactive interference compared with its uninduced control group. n = 8. Statistics: two-way ANOVA with Bonferroni’s multiple comparisons test. Results with error bars are means ± SEM. *p < 0.05. n.s., non-significant.

**Figure 4-figure supplement 1.**
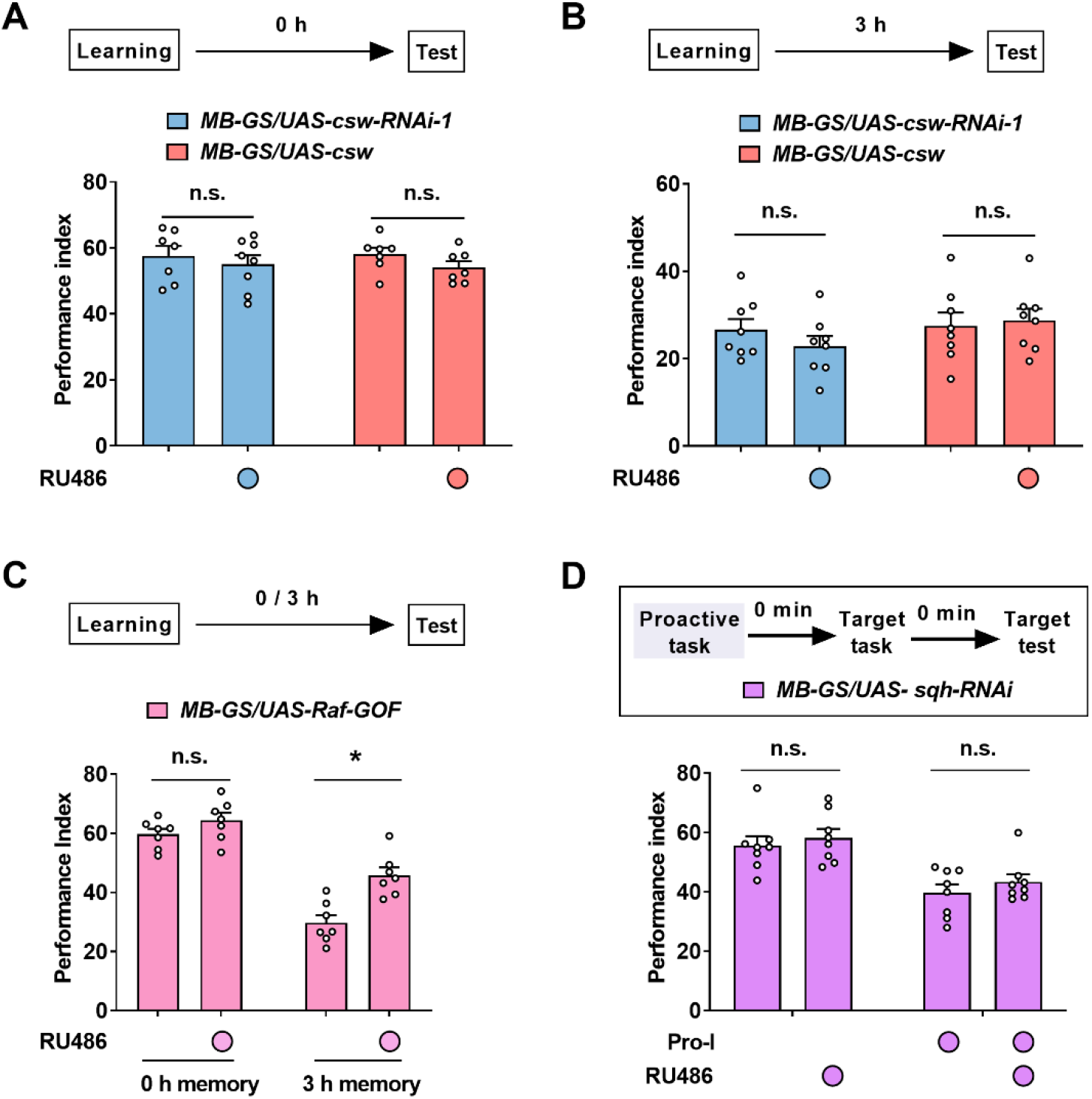
Behaviral data of a single learning task and proactive interference. (A) Learning performance. Acute knockdown or overexpression of CSW in MB neurons did not affect learning. n = 7-8. (B) 3-h memory performance. Acute knockdown or overexpression of CSW in MB neurons did not affect 3-h memory performance without proactive interference. n = 8. (C) Learning and 3-h memory performance. Acutely overexpressing Raf-GOF increased 3-h memory performance without affecting learning. n = 7. (D) The paradigm (top panel) and the behavioral results (bottom panel) of the proactive interference (Pro-I) experiment. Acute knockdown of Sqh in MB neurons (*MB-GS/UAS-sqh-RNAi*, RU486+) did not affect proactive interference. n = 8. Statistics: two-way ANOVA with Bonferroni’s multiple comparisons test. Results with error bars are means ± SEM. *p < 0.05. n.s., non-significant.

